# Molecular epidemiology of carbapenem-resistant *Enterobacter cloacae* complex infections uncovers high frequency of non-carbapenemase-producers in five tertiary care hospitals from Colombia

**DOI:** 10.1101/494807

**Authors:** Astrid V. Cienfuegos-Gallet, Ana M. Ocampo, Kalyan Chavda, Liang Chen, Barry N. Kreiswirth, J. Natalia Jiménez

## Abstract

**Background:** Infections caused by carbapenem-resistant *Enterobacter cloacae* (CR-Ecl) have been increasingly reported in the clinical setting; here we describe the clinical and molecular characteristics of CR-Ecl infections in a KPC endemic region.

**Methods:** A cross-sectional study was conducted in five tertiary-care hospitals in Medellín-Colombia. All patients infected by CR-Ecl from June-2012 to June-2014 were included. Sociodemographics and clinical information was retrieved from medical records. Antimicrobial susceptibility testing, phenotypic and molecular carbapenemase detection were performed. Analysis of *hsp60* and PFGE was done in a subset of isolates.

**Results:** Of 109 patients enrolled, 60.55% (66/109) were infected with non-carbapenemase-producing-Ecl (non-CP-Ecl). CP-Ecl patients were frequently hospitalized in the ICU (37.21% vs 12.12%) and had exposure to carbapenems (39.53% vs 15.15%) compared to non-CP-Ecl infected patients. All-cause 30-day mortality was higher in CP-Ecl than non-CP-Ecl infected patients (27.91% vs 19.70%). CP-Ecl harbored KPC-2 (83.72%) and KPC-3 (6.97%). Analysis of *hsp60* showed that CP-Ecl belonged primarily to cluster-VI of *Enterobacter xiangfangensis* (12/34) and cluster-XI (12/34) corresponding to *E. cloacae* subsp. *cloacae.* Non-CP-Ecl isolates belonged to cluster-VII/VIII (45/54), of *E. hormaechi* subsp. *steigerwaltii*. PFGE revealed isolates in cluster VII/VIII and XI were closely related within their own clusters.

**Conclusions:** The results revealed a high frequency of non-CP-Ecl among the CR-Ecl infections in a KPC endemic region, displaying distinct clinical and molecular characteristics in comparison to CP-Ecl. The study highlights a significant contribution of non-CP-Ecl to the prevalence of CR-Ecl. Infection control measures to curtail dissemination of CR-Ecl should not only focus on CP-Ecl but should also include non-CP-Ecl.

## Introduction

*Enterobacter* spp. have become a significant pathogen in clinical settings in the last decades (1). In fact, *Enterobacter cloacae* is among the top-five bacteria causing intra-abdominal infections in hospital and community settings in all regions of the world (2). Data from the National Nosocomial Infections Surveillance (NNIS) System from 1986–2003 in United States showed *Enterobacter* was a frequent cause of pneumonia (10.0%), bloodstream (4.4%), surgical site (9.0%) and urinary tract infections (UTI) (6.9%) among Gram-negative bacilli (3) and it was mainly associated with infections occurring in intensive care units (ICU) (4). Similarly, in Latin America, *Enterobacter* spp. is also among the top-five Gram-negative pathogens causing bloodstream infections (4.5%), pneumonia (5.1%) and skin and soft tissue infections (6.8%) (5).

*Enterobacter* spp. is increasingly associated with multi-drug resistance, including the resistance to the “last-resort” carbapenems. Wilson *et al.* (6) recently described two epidemics of carbapenem-resistant bacteria in United States. The first and most notorious started in 2000 and it was caused by the expansion of KPC-*K. pneumoniae* from the east to the pacific coast of the country, although resistance rates are decreasing in recent years. The second is the unfolding epidemic caused by carbapenem-resistant *E. cloacae* (CR-Ecl) extending from the east to the southwest and pacific coast of United States; in contrast to *K. pneumoniae*, rates of carbapenem resistance appear to be growing in *E. cloacae* in recent years. Additionally, multiple outbreaks of CR-Ecl harboring KPC (7,8), VIM (9), IMP (10) and OXA-48 (11) carbapenemases have been reported globally, highlighting the key role of *Enterobacter* in dissemination of carbapenem resistance.

In Colombia, *E. cloacae* is reported to be one of the most commonly isolated pathogens in both ICU and non-ICU wards (12). More alarmingly, the national surveillance data also showed that carbapenem-resistance rates in *E. cloacae* (10-16%) are similar to *K. pneumoniae* (14-15%) (12). However, KPC carbapenemases were only detected in 66% of *E. cloacae* isolates from the national program on antimicrobial resistance during 2012 to 2015, suggesting additional mechanisms mediating carbapenem-resistance in CR-Ecl (13). Despite the clinical importance of this pathogen, only a few studies have focused on the characterization of infections caused by CR-Ecl. The aim of this study is to describe the clinical and microbiological characteristics of infections caused by CR-Ecl in a KPC endemic region.

## Materials and Methods

### Study design and setting

A cross-sectional study was conducted in the city of Medellin, the second largest city in Colombia with 2.5 million inhabitants. The study was conducted in five tertiary care hospitals capturing both adult and pediatric populations. Hospital A (700-beds) and Hospital C (754-beds) are large university hospitals, Hospital B (286-beds) and Hospital D (300-beds) are medium size tertiary care centers and Hospital E (140-beds) is a small hospital specialized in cardiovascular diseases.

### Participants

All patients infected with carbapenem-resistant *Enterobacter cloacae* complex isolates in the five tertiary care hospitals from June 2012 to June 2014 were enrolled. Patients from any age, service and type of infection were included in the study at the time of the first CR-Ecl infection. Specialists in infectious diseases established the infection/colonization status of the patients using previous standardized definitions (14). The study protocol was approved by the Committee of Bioethics for Human Research at the Universidad de Antioquia (CBE-SIU) (approval no. 11-35-415) and by the Committee of Ethics at each of the participant institutions.

### Clinical information

Information retrieved from the medical records included sociodemographics (age and sex) and the following clinical variables: time at risk (defined as the number of days from admission to the date of sampling), transfer from another facility, ICU stay, use of invasive devices at the time of sampling or 48 hours before sampling, previous healthcare exposures (surgery in the previous year, prior ICU stay and antibiotic use in the last six months), dialysis, neutropenia, immunosuppressive conditions, comorbidities (trauma, cancer, diabetes mellitus, cystic fibrosis, neurological disease, cardiovascular disease, lung disease, burns, transplant, chronic obstructive disease), health care associated infection, mixed infection, empirical and targeted treatment, discharge (death, clinical improvement, cure). All clinical information was retrieved from the medical records and included in a standardized formulary by specialists in infectious diseases at each institution.

### Microbial identification and antimicrobial susceptibility testing

Identification of isolates and antimicrobial susceptibility testing was carried out by the Vitek® 2 automated system (bioMérieux, Marcy l’Etoile, France). Antibiotics tested included ceftriaxone, ceftazidime, cefepime, ertapenem, imipenem, meropenem, ciprofloxacin, gentamicin, amikacin, tigecycline and colistin. Resistant, intermediate or susceptible categories were defined following CLSI guidelines (15). Isolates were considered resistant to carbapenems if at least one of the carbapenems was non-susceptible (imipenem, meropenem or doripenem MIC ≥ 2 µg/mL or ertapenem MIC ≥ 1 µg/mL) (15).

### Phenotypic and molecular detection of β-lactamases

The three-dimensional test, which uses a mechanical lysate of the tested isolate to increase the sensitivity of carbapenemase detection (16), was performed in all CR-Ecl isolates. In addition, modified Hodge Test (MHT) (15) was conducted in a subset of 65 isolates. Molecular detection of *bla*_KPC_ variants were done using a molecular beacon-based real-time PCR assay (17). Isoforms of Tn*4401* element were evaluated by PCR (18). Detection of additional carbapenemase genes *bla*_VIM_, *bla*_IMP,_ *bla*_NDM_ and *bla*_OXA-48_ were done by conventional multiplex PCR (19). In addition, extended spectrum *β*-lactamases (ESBLs) genes *bla*_CTX-M-1_, *bla*_CTX-M-2_, *bla*_CTX-M-_ 8, *bla*_CTX-M-9_, *bla*_CTX-M-25_, *bla*_TEM_ and *bla*_SHV_ were evaluated using PCR and Sanger sequencing (20), and acquired AmpC genes *bla*_ACT/MIR_, *bla*_CMY-1/MOX_, *bla*_CMY-2/LAT_, *bla*_FOX_, *bla*_DHA_ and *bla*_ACC_ were assessed using PCR (20).

### Sequence analysis of *ompF*

To investigate additional mechanisms related to carbapenem resistance in *E. cloacae* isolates, the full-length sequences of *ompF* were analyzed in a subset of 91 isolates. Sequences were compared to the reference strain *E. cloacae* NCTC 13405 (KT780421). All sequences were translated and aligned using MUSCLE accessory application in Geneious® 8.1.9 (21).

### *hsp60* phylogenetic analysis

Sequences of *hsp60* were obtained from 91 CR-Ecl isolates using primers and conditions previously reported (22). Isolates were classified according to the Hoffman and Roggenkamp scheme (22), by comparison with reference strains of *E. cloacae* complex retrieved from GenBank. Sequences sets were aligned using MUSCLE accessory application in Geneious® 8.1.9 (21). Bayesian phylogenetic analysis was performed using Markov chain Monte Carlo (MCMC) sampling implemented in MrBayes 3.2.6 (23), under a TPM3+G model selected according to the Bayesian Information Criteria in jmodeltest2 (24). The MCMC search was run for 3 x 10^6^ generations with trees sampled every 500^th^ generations and burn-in length of 200.000. Parameters estimates were assessed in Tracer v1.6 (available at http://tree.bio.ed.ac.uk/software/tracer/). *hsp60* sequence of *E. aerogenes* NBRC13534 (AB375469) was used as an outgroup.

### Strain genotyping

Genotyping of isolates was performed by pulsed-field gel electrophoresis (PFGE) on 68 randomly selected isolates including isolates from each hospital. PFGE conditions were described previously (25). Briefly, DNA was digested with 20U of *Xba*I restriction endonuclease at 37°C for two hours. PFGE conditions were initial switch time 2.2 sec, final switch time 63.8 sec, angle 120° and 6 v/cm volts. PFGE was run for 24 hours. Analysis of relatedness among *E. cloacae* complex isolates was performed on BioNumerics software version 6.0 (Applied Maths, Sint-Martens-Latem, Belgium) using the Dice coefficient and a cutoff value of 80 or higher for genetic relatedness. For dendogram generation the unweighted-pair group method analysis with average linkages (UPGMA) was used. DNA fragment patterns were normalized using a bacteriophage lambda ladder PFGE marker (New England BioLabs, United Kingdom) with a 1% position tolerance for further analysis. Six isolates from current study were characterized by WGS analysis in a previous report (26). Isolates sequenced were: EL012 (44541, accession no: NZ_JZXU01000000), EL036 (44565, NZ_JZXT01000000), EH005 (44517, NZ_JZXX01000000), EH012 (44524, NZ_JZXW01000000), EH015 (44527, NZ_JZXV01000000) and EP004 (44589, NZ_JZXS01000000).

### Accession numbers

All of the *hsp60* sequences determined in this study are available at NCBI under accession numbers EL044 (MH614175), EH006 (MH614176), EH019 (MH614177), EP010 (MH614178), EL033 (MH614179), EP007 (MH614180), EH010 (MH614181), ER005 (MH614182), EH011 (MH614183), EL018 (MH614184), EP008 (MH614185), EP005 (MH614186), EC001 (MH614187), EH015 (MH614188), ER002 (MH614189), EL036 (MH614190), EL021 (MH614191), EH004 (MH614194), EC003 (MH614195), EH005 (MH614197).

### Statistical analysis

To describe patient’s characteristics, absolute and relative frequencies were used for qualitative variables, and median and interquartile range for quantitative variables with non-normal distribution. All statistical analyses were performed in STATA/IC 15.1.

## Results

### Description of patients infected with carbapenem-resistant *E. cloacae* complex

A total 109 patients were infected with CR-Ecl during the study period. The most frequent infections in the study population were surgical site infections (n=25, 22.94%), followed by intra-abdominal (n=18, 16.51%), and urinary tract infections (UTI) (n=18, 16.51%). Overall all-cause in-hospital mortality was 24.77% (n=27) and all-cause 30-day mortality was 22.94% (n=25) (Table 1). The majority of patients were older (median age 64 years, IQR 49 – 74) and from male (n=73, 66.97%). Almost all infections in the study population were healthcare-associated (n=108, 99.08%). CR-Ecl frequently infected patients with comorbidities (92.66%) and with at least one medical device at the time of sampling or 48 hours before (n=70, 64.22%), mainly urinary catheters (n=50, 45.87%). Most CR-Ecl infected patients had previous antibiotics use in the last six months (n=90, 84.11%), mostly piperacillin/tazobactam (n=51, 46.79%), fluoroquinolones (n=29, 26.61%) and carbapenems (n=17, 24.77%). Patients also have history of hospitalization in the last six months (n=82, 75.23%) and surgery in the last year (n=76, 69.72%). Additional patient’s characteristics are presented in Table 1.

**Table 1.**
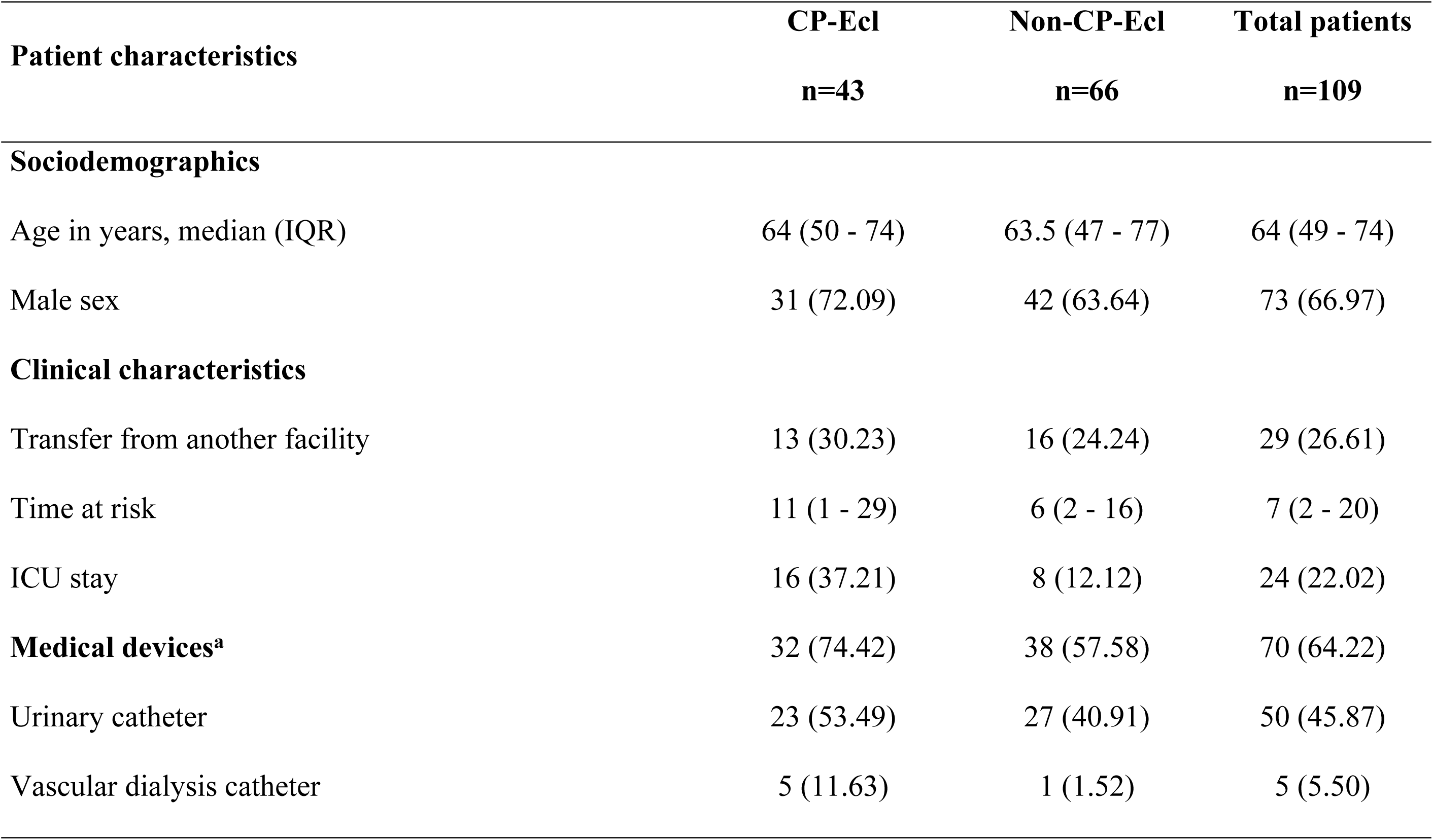
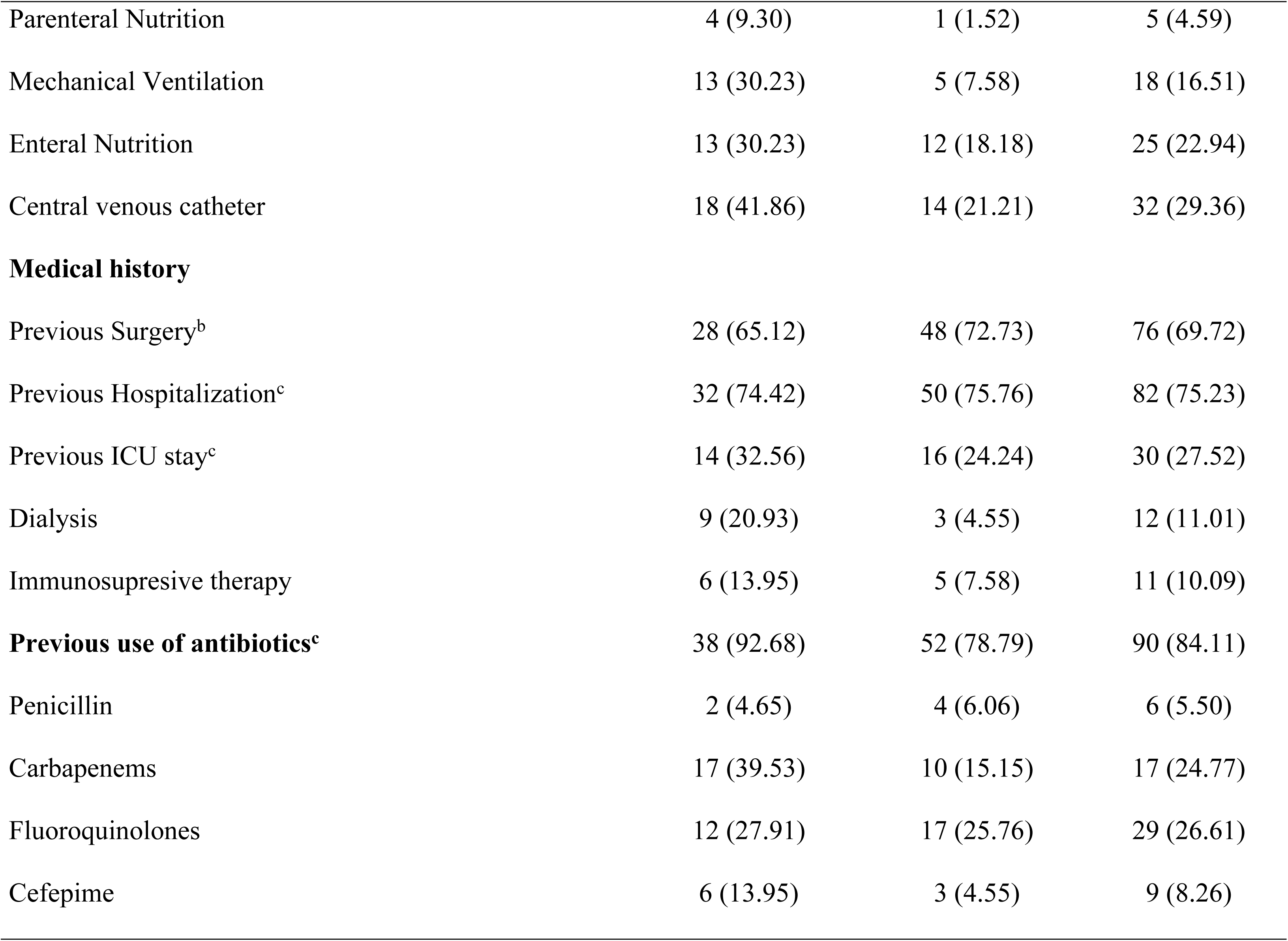
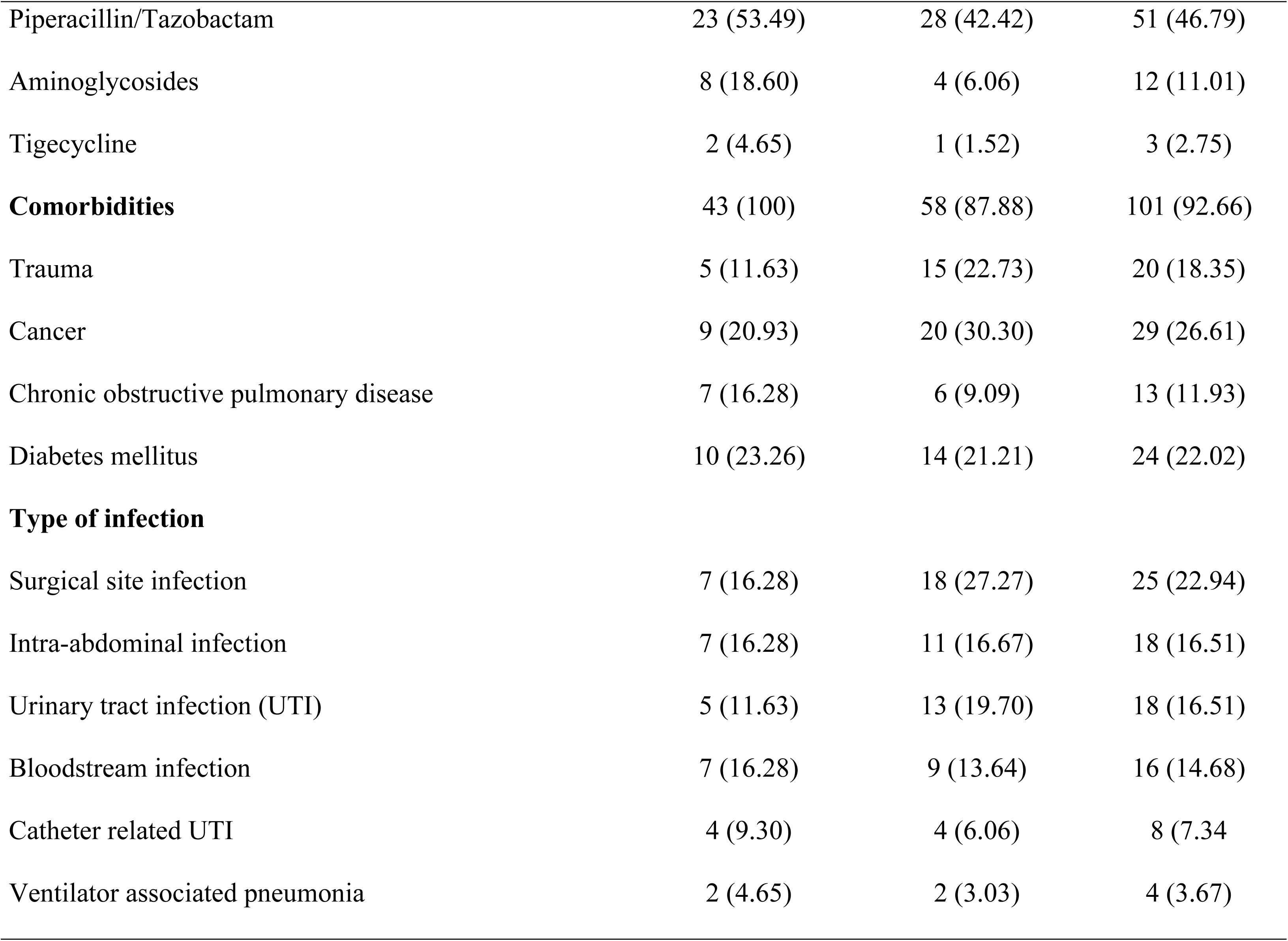
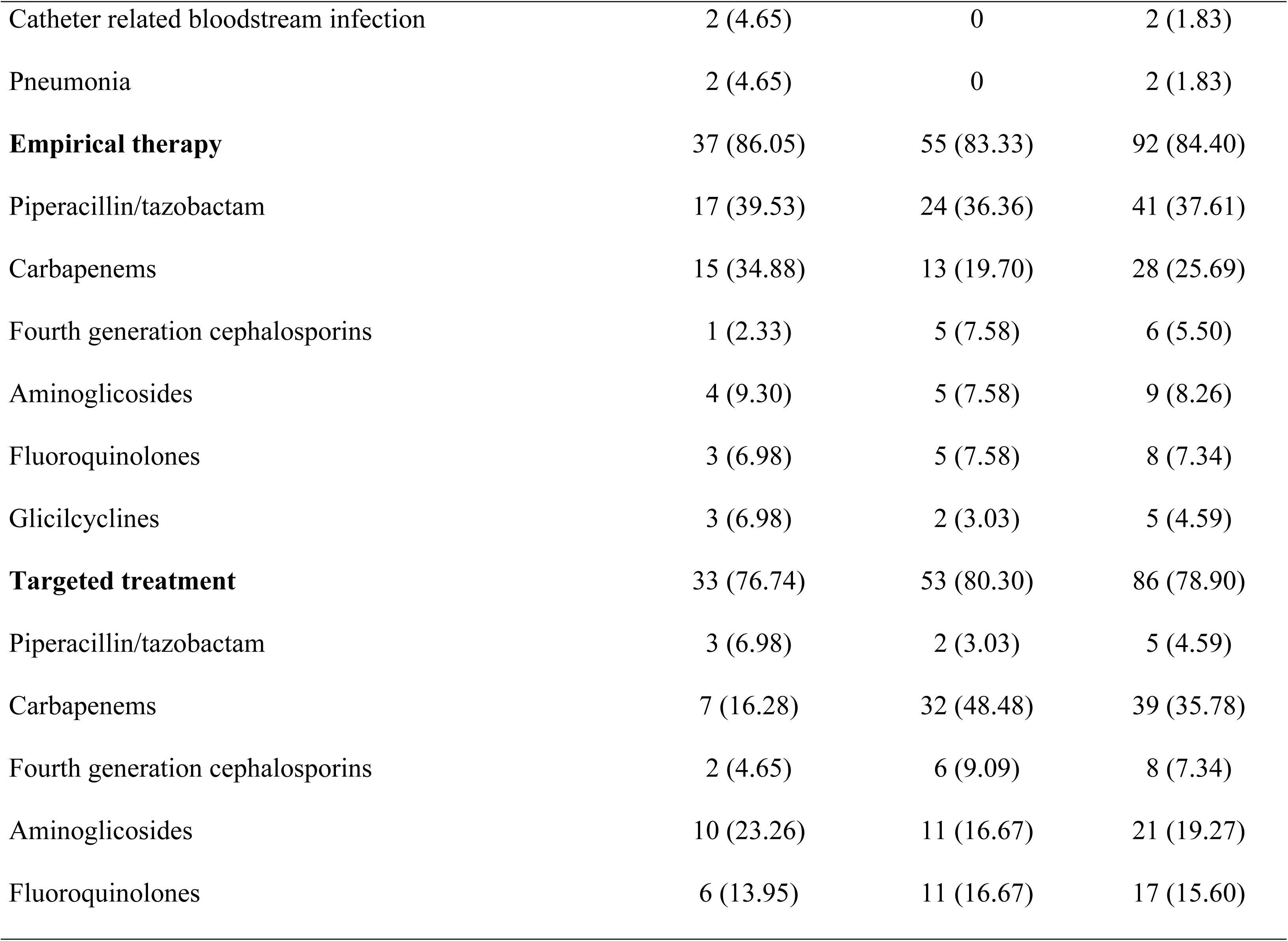
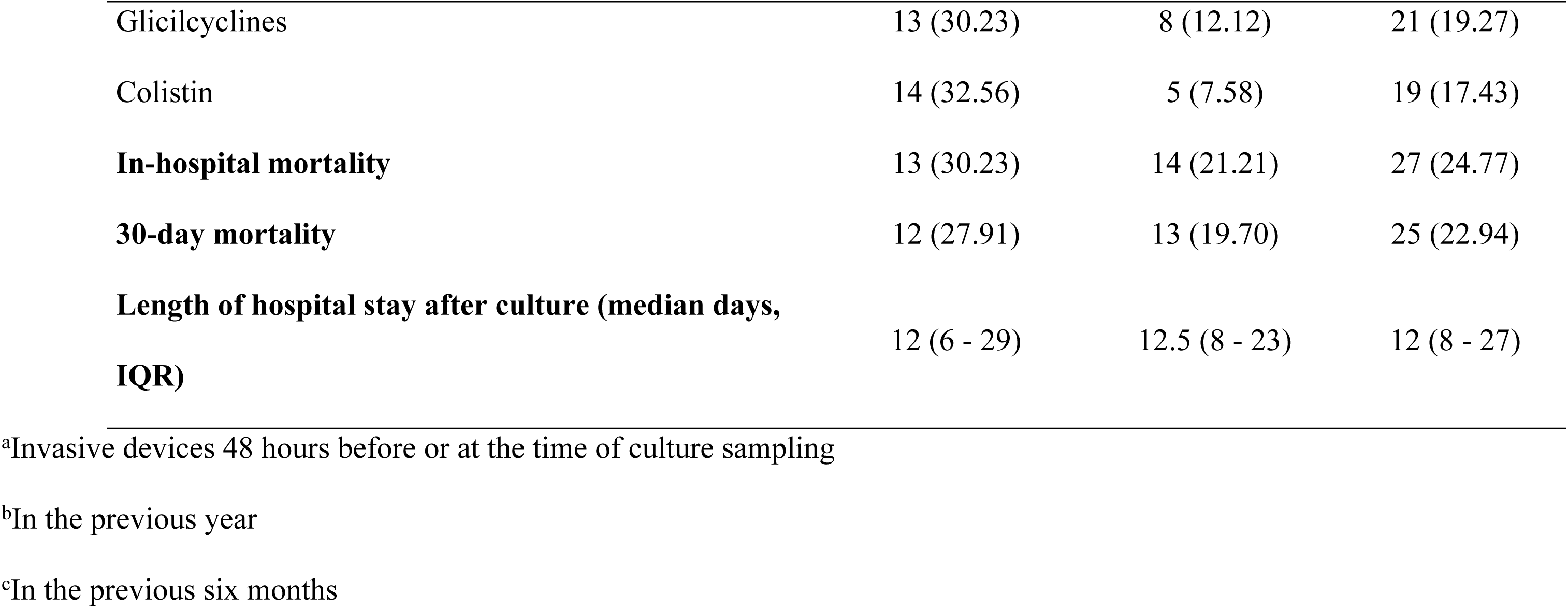
Clinical characteristics of patients infected with carbapenem-resistant *Enterobacter cloacae* complex according KPC detection.

### Phenotypic and molecular detection of carbapenemases

43 strains were found to harbor *bla*_KPC_, including *bla*_KPC-2_ (83.72%) and *bla*_KPC-3_ (6.97%), with most of them in the Tn*4401* isoform b (41/43); one isolate carried Tn*4401* isoform a and the remaining isolate was negative for Tn*4401*. No other carbapenemase genes (*bla*_VIM_, *bla*_IMP_, *bla*_NDM_ and *bla*_OXA-48_) were detected. Notably, a higher frequency of non-carbapenemase-producing *E. cloacae* (non-CP-Ecl) (66/109, 60.55%) were detected among CR-Ecl. In addition, a high rate of false-positive results were observed in the three-dimensional test (n=104, 95.41%), although only 39.55% of isolates harbored *bla*_KPC_. Similarly, MHT was positive in 38/65 (58.46%) isolates, but only 19 (50%) harbored *bla*_KPC_.

### Clinical and microbiological description of infections caused by non-carbapenemase-producing *E. cloacae*

Patients infected with non-CP-Ecl presented mainly with surgical site infections (n=18, 27.27%), followed by urinary tract infections (n=13, 19.70%) and intra-abdominal infections (n=11, 19.70%). In hospital mortality was 21.21% (n=14) and length of hospital stay after positive culture was 12.5 days (IQR 8-23). Patients were treated frequently with monotherapy (n=32, 48.48%), mostly with carbapenems (18/32) or fluoroquinolones (5/32). Combined therapy was administered in 27.27% (n=18) of patients, mainly carbapenems plus aminoglycosides (4/18), glicilcyclines (3/18) and polymixins (3/18). Additional characteristics of patients are presented in Table 1.

Non-CP-Ecl frequently harbored chromosomal β-lactamases gene *bla*_ACT/MIR_ (n=32, 53.33%) and *bla*_ACT/MIR_+*bla*_TEM-1_ (n=11, 18.33%). Interestingly, additional analysis of *ompF* revealed 37/66 isolates had premature stop codons: D166X (20/37), E65X (10/37), F145X (4/37), K3X (1/37), Q93X (1/37) and L105X (1/37). Meanwhile, 17/66 isolates had frameshift mutations and 23/66 had missense mutations (V141A and V141K) in loop 3, which constitutes the eyelet of the porin channel. In total 55/66 isolates had at least one of the above described mutations in the *ompF* sequence.

Of note, non-CP-Ecl exhibited high frequency of ertapenem resistance (85.94%), but lower resistance to other carbapenems (imipenem 19.35% and meropenem 3.03%). Resistance to gentamicin, ciprofloxacin and tigecycline were also frequently observed in non-CP-Ecl, but most isolates were susceptible to colistin (96.23%) (Table 2). The antibiotic resistance profile showed that most non-CP-Ecl were resistant to ertapenem+amikacina+gentamicin+ciprofloxacin (n=21, 31.82%).

**Table 2.** Microbiological characteristics of carbapenem-resistant *Enterobacter cloacae* complex isolates.

| Isolate characterization | CP-Ecl<br>n=43 | Non-CP-Ecl<br>n=66 | Total isolates<br>n=109 |
| --- | --- | --- | --- |
| Threedimensional test | 43 (100) | 61 (92.42) | 104 (95.41) |
| Antimicrobial resistance pattern |  |  |  |
| Ceftazidime | 37 (92.50) | 47 (78.33) | 84 (84.00) |
| Ceftriaxone | 33 (94.29) | 42 (84.00) | 75 (88.24) |
| Cefepime | 27 (65.85) | 13 (20.00) | 40 (37.74) |
| Ertapenem | 33 (91.67) | 55 (85.94) | 88 (88.00) |
| Imipenem | 40 (93.02) | 12 (19.35) | 52 (49.52) |
| Meropenem | 38 (88.37) | 2 (3.03) | 40 (36.70) |
| Gentamicin | 26 (60.47) | 39 (60.00) | 65 (60.19) |
| Amikacin | 21 (48.84) | 8 (12.12) | 29 (26.61) |
| Ciprofloxacin | 27 (62.79) | 44 (66.67) | 71 (65.14) |
| Tigecycline | 29 (78.38) | 36 (75.00) | 65 (76.47) |
| Colistin | 2 (6.25) | 2 (3.77) | 4 (4.71) |
| Multidrug resistant | 21 (70.00) | 33 (73.33) | 54 (72.00) |

Bayesian analysis of *hsp60* sequences of 54 out of 66 isolates non-CP-Ecl strains revealed most isolates belonged to cluster VII/VIII (45/54), followed by cluster VI (4/54) (Figure 1 and 3). Previous phylogenomics analysis, including several isolates from this study (EH005, EH012, EL012 and EP004), assigned cluster VII/VIII strains as *E. hormaechi* subsp. *steigerwaltii*. Only eight (8/53, 15.1%) isolates from this cluster carried KPC-2. Of note, 44/45 non-CP-Ecl isolates from this cluster had the following mutations in *ompF*: premature stop codons, mainly D166X (20/45), E65X (10/45) and F145X (4/45), frameshifts in the protein sequence (9/45) and alterations in the loop3 of the porin channel (V141A, 10/45), suggesting the carbapenem resistance in this cluster may be associated with the OmpF dysfunction. Additionally, PFGE analysis of 34 isolates from cluster VII/VIII revealed 12 isolates were closely related (Dice’s coefficient >80%) (Figure 2, Table S1).

**Figure 1.**
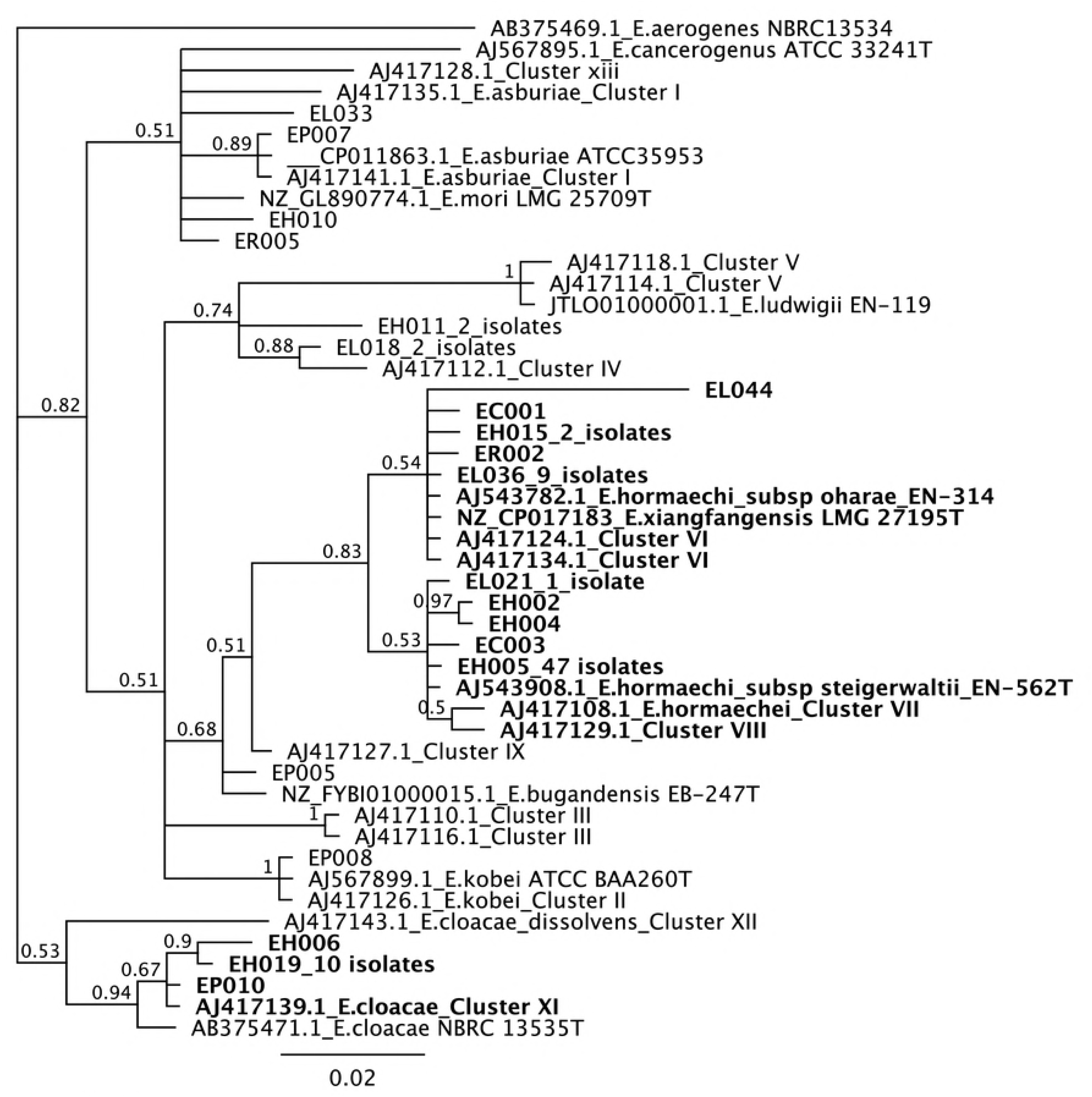
Bayesian phylogenetic tree depicting the relationship among *E. cloacae* complex isolates based on *hsp60* sequences obtained from 91 study isolates and 27 *hsp60* sequences of reference strains retrieved from GenBank. The Bayesian tree was constructed using the TMP3+I+G nucleotide substitution model. The strain *E. aerogenes* NBRC13534 was used as outgroup. Posterior probabilities are shown in each node. A high posterior probability support (>80%) was found for the clades with isolates from cluster I, cluster II, cluster IV, cluster VI/VII/VIII and cluster XI. A low posterior probability support was found for clades with *E. hormaechi* subspecies (54 and 53%). Isolates belonging to the three largest clusters are shown in bold.

**Figure 2.**
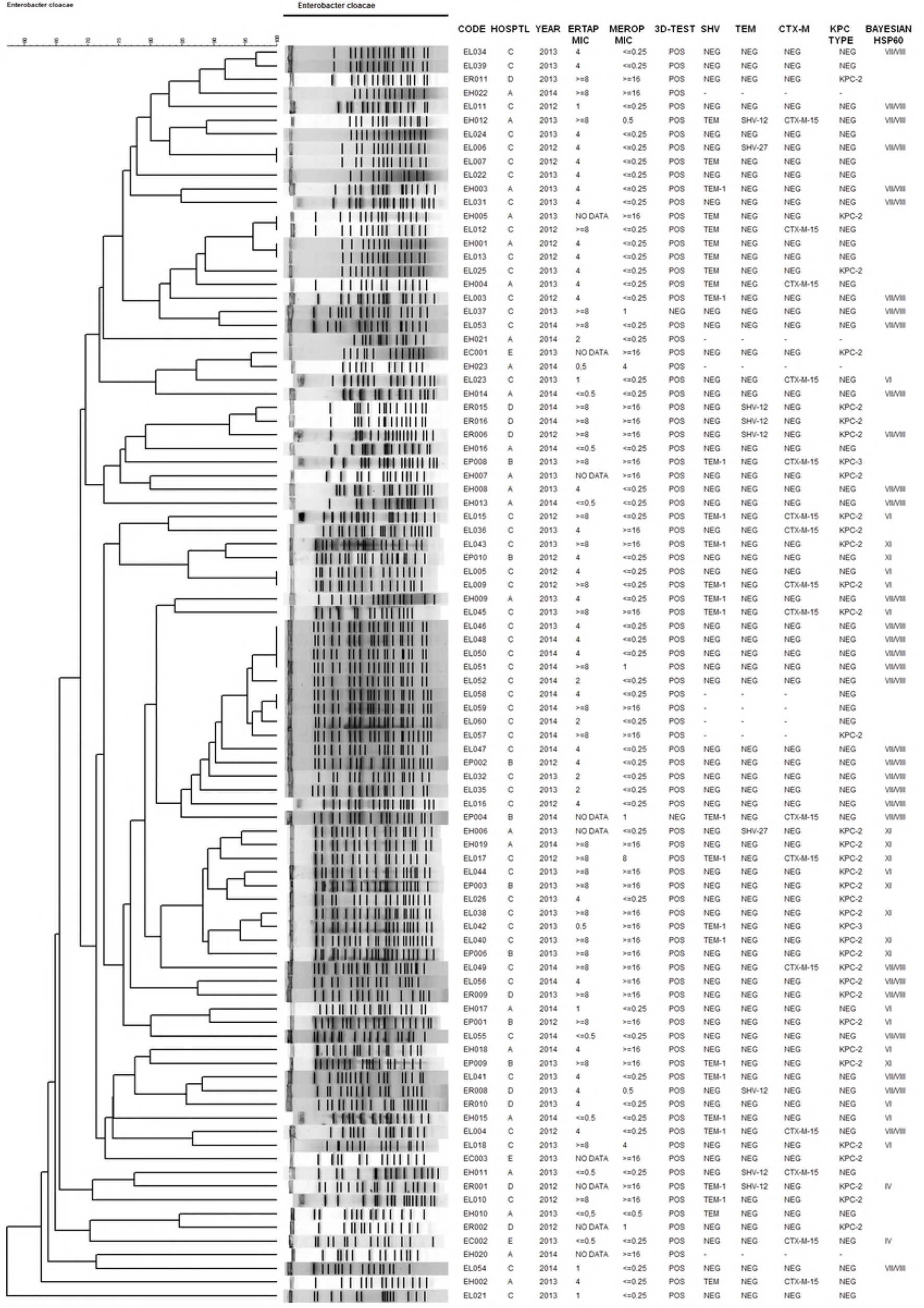
PFGE dendrogram showing the genetic relationship among 68 isolates of carbapenem-resistant *E. cloacae* complex. The Dice similarity coefficient and the unweighted pair group method with arithmetic averages were used for dendogram generation in Bionumerics software. Two groups of isolates were closely related according to the Dice coefficient (>80%); the first group were mainly KPC-negative and correspond mostly to cluster VII/VIII by *hsp60* phylogenetic analysis, the second group are mainly KPC-positive, and correspond primarily to cluster XI by *hsp60* analysis.

Among non-CP-Ecl, Hospital C accounted for 32/45 of cluster VII/VIII isolates, followed by Hospital A (9/45). PFGE pulsotypes were closely related among non-CP-Ecl isolates from Hospital C, suggesting a clonal dissemination of the similar non-CP-Ecl in this hospital (Figure 2 and 3). The majority of isolates from cluster VII/VIII were recovered spanning the two-year study period (Figure S2). Isolates from this cluster were found in a variety of infections (Figure S1).

Figure 3. (A) Distribution of *hsp60* clusters among KPC and non-KPC-Ecl isolates. (B) PFGE pulsotypes identified within cluster XI (CP-Ecl) and (C) cluster VII/VIII (non-CP-Ecl) isolates.

### Clinical and microbiological description of infections caused by carbapenemase-producing *E. cloacae*

Patients infected with CP-Ecl presented diverse infections including surgical site (n=7, 16.28%), intra-abdominal (n=7, 16.28%), primary bloodstream (n=7, 16.28%) and urinary tract infections (n=5, 11.63%). The targeted treatment of CP-Ecl infected patients was predominantly monotherapy (n=16, 37.21%), with tigecycline (4/16) or polymixins (5/16).

Regarding microbiological characteristics, CP-Ecl frequently carried additional β-lactamases, such as *bla*_ACT/MIR_, *bla*_CTX-M-15_, *bla*_TEM-1_ and *bla*_SHV-12_. The most frequent β-lactamase profile was *bla*_ACT/MIR_+*bla*_KPC-2_ (n=10, 25.00%).

Susceptibility testing showed CP-Ecl isolates had high frequency of resistance to different antibiotics, including cefepime (65.85%), imipenem (93.02%), meropenem (88.37%) and amikacin (48.84%), but were susceptible to colistin with a MIC ≤ 0.5 μg/mL (93.75%) (Table 2). The most common profile in KPC-Ecl was resistance to all antibiotics except colistin.

Bayesian phylogenetic analysis of *hsp60* from 34 CP-Ecl isolates revealed most strains belonged primarily to cluster VI (12/34) and XI (12/34) (Figure 1 and 3). Cluster VI includes *E. hormaechi* subsp. *oharae* and the recently described *E. xiangfangensis.* Previous phylogenomics analysis of two isolates (EH015 and EL036) from this cluster identified them as *E. xiangfangensis* (26). Most isolates of cluster VI harbored KPC-2 (11/12) and were genetically diverse according PFGE (Figure 1 and 3).

Remarkably, 10/12 cluster XI isolates were closely related according PFGE analysis (Dice coefficient >80%) (Figure 2, Table S2), 11/12 harbored KPC-2 and all harbored the same insertion in *ompF* (VT in pos. 45) and the premature stop codon (L105X). Most KPC-positive infected patients were from Hospital C (17/34), from which seven belonged to cluster XI. Cluster XI isolates caused mainly UTI and intra-abdominal infections (Figure S1).

### Comparison of clinical characteristics between patients infected by carbapenemase-producing and non-carbapenemase producing *Enterobacter cloacae* complex

Patients infected with CP-Ecl and non-CP-Ecl were identified along the two years of the study with no cluster of cases over time (Figure S2). CP-Ecl patients were frequently hospitalized in the ICU (37.21% vs 12.12%) and had at least one medical device at the time or 48 hours before sampling, such as mechanical ventilation (30.23% vs 7.58%) and central venous catheter (41.86% vs. 21.21%) In addition, CP-Ecl infected patients had frequent exposure to carbapenems (39.53%) compared to non-CP-Ecl infected patients (15.15%).

All-cause in-hospital mortality was higher among CP-Ecl (30.23%) than in non-CP-Ecl (21.21%) among infected patients. Similar findings were observed for all-cause 30-day mortality (27.91% in CP-Ecl and 19.70% in non-CP-Ecl). The median time to death was 6 days (IQR 3-11) in CP-Ecl and 9 days (IQR 3-14) in non-CP-Ecl infected patients.

## Discussion

Colombia is regarded as one of the KPC endemic regions, and approximately 80 to 86% of carbapenem-resistant isolates carried KPC carbapenemases, mostly in *K. pneumoniae* (12,27). Limited reports about CR-Ecl were available in the country. The national surveillance system reported in 2015 that most (66.1%) carbapenem-non-susceptible *E. cloacae* isolates carried KPC (144/218), followed by KPC+GES (7/218), VIM (5/218), NDM (4/218), KPC+VIM (2/218) and GES (1/218). By contrast, 25.22% (55/218) of carbapenem-non-susceptible *E. cloacae* did not harbor any carbapenemase (12). A similar study conducted in eight Colombian regions showed that most CR-Ecl isolates (19/28, 67.85%) harbored KPC (28). By contrast, our study uncovered a significantly higher proportion (60.55%) of non-CP-Ecl in CR-Ecl.

In our study a higher proportion of isolates (60.55%), did not harbor carbapenemases (KPC, NDM, VIM, IMP or OXA-48), and remarkably, *hsp60* phylogenetic analysis revealed the predominance of cluster VII/VIII in non-CP-Ecl isolates. In addition, PFGE showed a predominant pulsotype within the non-CP-Ecl cluster VII/VIII, suggesting a clonal spread of the same strain in at least 12 patients. Previous phylogenomics analysis using core SNPs and average nucleotide identity (ANI) confirmed identification of four isolates from this cluster as *E. hormaechi* subsp. *steigerwaltii*. In contrast to previous reports reporting genetic diversity among non-CP-Ecl (29), our findings demonstrated the clonal dissemination of non-CP-Ecl strains.

Carbapenem resistance in non-CP-Ecl isolates could be the result of β-lactamase production and porin loss. Pecora *et al.* (30), found that 32.4 to 52.3% of *E. cloacae* isolates not carrying carbapenemases genes but β-lactamase genes (CTX-M-15, TEM-116, AmpC) in conjunction with OmpC and OmpF defects, were resistant to carbapenems. This agreed with our results where a high rate of false-positives in the MHT and three-dimensional test was detected in non-CP-Ecl isolates. In fact, Wang *et al.* (31) reported 3.3% of false positive in non-carbapenemase producing but ESBL-producing Enterobacteriaceae, mainly CTX-M producers, indicating the lack of specificity of MHT for detection of serine carbapenemases. In our study, CTX-M-15 was only detected in 8 out of 66 non-CP-Ecl. In addition, these isolates also carried SHV-12 and SHV-27 ESBLs (10/66) and several mutations and premature stop codons in *ompF*. It is also important to highlight that in our study only 2/66 of non-CP-Ecl and 38/43 CP-Ecl were resistant to meropenem, suggesting meropenem resistance may be useful for suspecting the presence of KPC in *E. cloacae* complex.

In our study, ∼40% of the isolates were carbapenemase-producers, with 83.72% harboring KPC-2 and 6.97% harboring KPC-3. Similarly, carbapenem resistance in *E. cloacae* is also associated with the presence of KPC in the USA, where outbreaks of closely related isolates of *E. cloacae* clone ST171 harboring KPC-3 have been described in Minnesota and in North Dakota (8,32). In our study, a small outbreak of CP-Ecl harboring KPC-2 occurring in one of the hospitals was also detected. *hsp60* phylogenetic analysis revealed among CP-Ecl the clusters VI, XI and VII/VIII of the Hoffman and Roggenkamp scheme (22), but the majority of isolates from cluster XI, which comprises *E. cloacae* subsp. *cloacae,* shared the same pulsotype, suggesting transmission of the CP-Ecl strain in at least ten patients. Cluster VI and VII/VIII was also predominant in CP-Ecl isolates, however pulsotypes were diverse within these groups. These results suggest that in addition to clonal spread, horizontal transfer of KPC plasmids in diverse genetic backgrounds are important mechanisms in dissemination of CP-Ecl in our setting.

Noteworthy, CP-Ecl and non-CP-Ecl displayed distinct clinical characteristics and resistance patterns. Overall, non-CP-Ecl infected patients were hospitalized in general wards, while CP-Ecl patients were frequently hospitalized in the ICU, had mechanical ventilation, central venous catheter, vascular dialysis catheter and previous carbapenem exposure. Importantly, observed all-cause mortality (30.23% vs 21.21%, respectively) and 30-day mortality (27.91% vs 19.70%, respectively) was higher in CP-Ecl group compared to non-CP-Ecl. Other studies have reported a comparable mortality rate in patients infected with KPC-Ecl, close to 35% (7/20), with 15% (3/20) of deaths attributable to CRE infections (7). It is important to highlight that ICU stay at the time of sampling and previous use of carbapenems were more frequent in the CP-Ecl infected group than in the non-CP-Ecl group. In support of these findings a previous study reported that ICU admission at the time of infection was common in CP-Ecl infections (7) and recently, Okamoto *et al.* (33) found that exposure to carbapenems (OR, 2.25; 95% CI, 1.06–4.77) was an independent risk factor for KPC-producing Gram-negative acquisition, in addition to colonization pressure (OR, 1.02; 95% CI, 1.01–1.04) and comorbidities measured by the Charlson index (OR, 1.14; 95% CI, 1.01–1.29). Although limited studies on CP-Ecl epidemiology are available, study addressing the identification of factors associated with ESBL enzymes in *E. cloacae* infections showed that the previous use of antibiotics (46/70 vs 17/20) in addition to mechanical ventilation (47/70 vs 19/20), were frequent in ESBL-positive-Ecl compared to ESBL-negative-Ecl infected patients, respectively (34). These results agreed with the preponderant role of carbapenems (33,35–37) and ICU stay (38–40) as risk factors for KPC-*K. pneumoniae* infection.

This study has limitations. First, description of clinical and microbiological variables such as the colonization status of CR-Ecl patients was not included because it was missing for 42.20% of patients included in the study. Similarly, results from the MHT were only described for 65 isolates with available information. In addition, ompC sequencing analysis were not included in this study, due to the high diversity of ompC sequences among different clusters.

In conclusion, our study revealed important differences in the molecular epidemiology of CR-Ecl in a KPC endemic setting, where non-CP-Ecl accounted for the majority of CR-Ecl infections. Both clonal spread and plasmid transfer are involved in CP-Ecl infections, while clonal spread was found the major mechanism in dissemination of non-CP-Ecl. Overall both CP-Ecl and non-CP-Ecl infections have similar clinical characteristics, but CP-Ecl infected patients had high mortality, were frequently hospitalized in ICU, have invasive devices and previous carbapenem exposure. The MHT and three-dimensional test showed false positives for carbapenemase detection, but resistance to meropenem instead of ertapenem could help in the detection of CP-Ecl. Finally, the study highlights significant contribution of non-CP-Ecl to the overall prevalence of CR-Ecl. Infection control measures to curtail dissemination of CR-Ecl should not only focus on CP-Ecl but also include non-CP-Ecl.

## Acknowledgment

This work was supported by Colciencias, Project code 111565741641 to J.N.J. This work was also supported by the following grants from the National Institute of Allergy and Infectious Diseases (R01AI090155 to B.N.K and R21AI117338 to L.C).

## Supporting Information

**Table S1.** Clinical characteristics of patients infected with non-CP-Ecl isolates closely related by PFGE (Dice’s coefficient >80) from cluster VII/VIII.

**Table S2.** Clinical characteristics of patients infected with CP-Ecl isolates closely related by PFGE (Dice’s coefficient >80) from cluster XI.

**Figure S1.** Type of infections caused by CP-Ecl and non-CP-Ecl according to hsp60 clusters. (A) CP-Ecl isolates and (B) Non-CP-Ecl isolates.

**Figure S2.** Epidemic curve of carbapenem-resistant E. cloacae complex infected patients (A) according E. cloacae genetic cluster of Hoffman & Roggenkamp in Hospital C and (B) according KPC and non-KPC harboring E. cloacae.

